# Optimization of Bio-Orthogonal Non-Canonical Amino acid Tagging (BONCAT) for effective low-disruption labelling of Arabidopsis proteins *in vivo*

**DOI:** 10.1101/2025.05.22.655591

**Authors:** Nicholas Hassan, Shelly Braun, Mohana Talasila, Curtis Kennedy, Luke Yaremko, Richard P. Fahlman, R. Glen Uhrig

## Abstract

Plants require robust responses in protein synthesis to adapt to variable environmental conditions. Measurement of newly synthesized proteins has been successfully facilitated with Bio-Orthogonal Non-Canonical Amino acid Tagging (BONCAT) across a multitude of organisms. Here, we use non-canonical amino acids (NCAAs) L-azidohomoalanine (AHA) or L-homopropargylglycine (HPG) incorporation in place of methionine residues into the actively translating *Arabidopsis* proteome, allowing for that subset of proteins to be enriched for mass spectrometry quantification. Although this technique has seen occasional use in plants, optimization of the protocol to maximize functionality while minimizing organismal stress has not yet been established. Here, we provide evidence for successful implementation through the liquid immersion of seedlings in AHA or HPG-containing media that functions with significantly lower concentrations than the literature standard. Our approach splits acute exposure and incorporation phase of labelling to mitigate potential negative impacts of prolonged NCAA exposure without compromising effective enrichment capacity, and demonstrate that this results in an unperturbed growth phenotype for AHA-treated seedlings. Finally, we demonstrate the capacity of this modified approach to enrich newly synthesized proteins from the whole proteome under standard stress conditions. These improvements allow for a broader use of BONCAT technologies in molecular plant research, affording a deeper understanding of the newly synthesized proteome without negatively impacting plant health.

## INTRODUCTION

Balancing protein synthesis and degradation defines proteostasis in all cells, which in turn defines the function of all organisms. Rapid adjustment of protein levels and proteome plasticity is essential for survival in a changing environment, especially for plants, which are carefully tuned to their environmental changes. Defining newly synthesized proteins, or the nascent proteome, is crucial to understanding plant cell regulation and adaptation to environmental stressors. However, conventional strategies for labelling the newly synthesized proteome, such as radiolabelling, lack depth of analysis (Tivendale et al., 2021), while characterization of the nascent proteome by enrichment-based mass spectrometry (MS) approaches can better quantify changes in lower abundant proteins. To fill this niche, Bio-Orthogonal Non-Canonical Amino acid Tagging (BONCAT), has been developed to label nascent proteins with a methionine analog, allowing for enrichment of newly synthesized proteins (Dieterich et al., 2006; Parker and Pratt, 2020).

BONCAT utilizes a combination of azide and alkyne groups to form irreversible conjugations via a copper-catalyzed azide-alkyne cycloaddition (CuAAC; reviewed in Chen et al., 2023). To employ this in a biological setting, unnatural methionine analogs L-azidohomoalanine (AHA, azide) or L-homopropargylglycine (HPG, alkyne) are supplied to live organisms. These non-canonical amino acids (NCAAs) are incorporated into proteins during synthesis by the relatively promiscuous methionine tRNA (Figure 1A; Bagert et al., 2014). Labelled proteins are then extracted and “clicked” to the opposing azide or alkyne conjugated to biotin, effectively biotinylating newly synthesized proteins to facilitate enrichment via streptavidin coupled matrix for subsequent quantification via liquid chromatography (LC)-MS/MS. While BONCAT was initially deployed in mammalian cells, it has since found use in a wide variety of organisms, such as live mice, bacteria, and archaea (Dieterich et al., 2006; Kern and Ferreira-Cerca, 2022; Lindivat et al., 2021; Saleh et al., 2019). However, unlike other organisms, BONCAT implementation in plants has not been fully realized.

**Figure 1.**
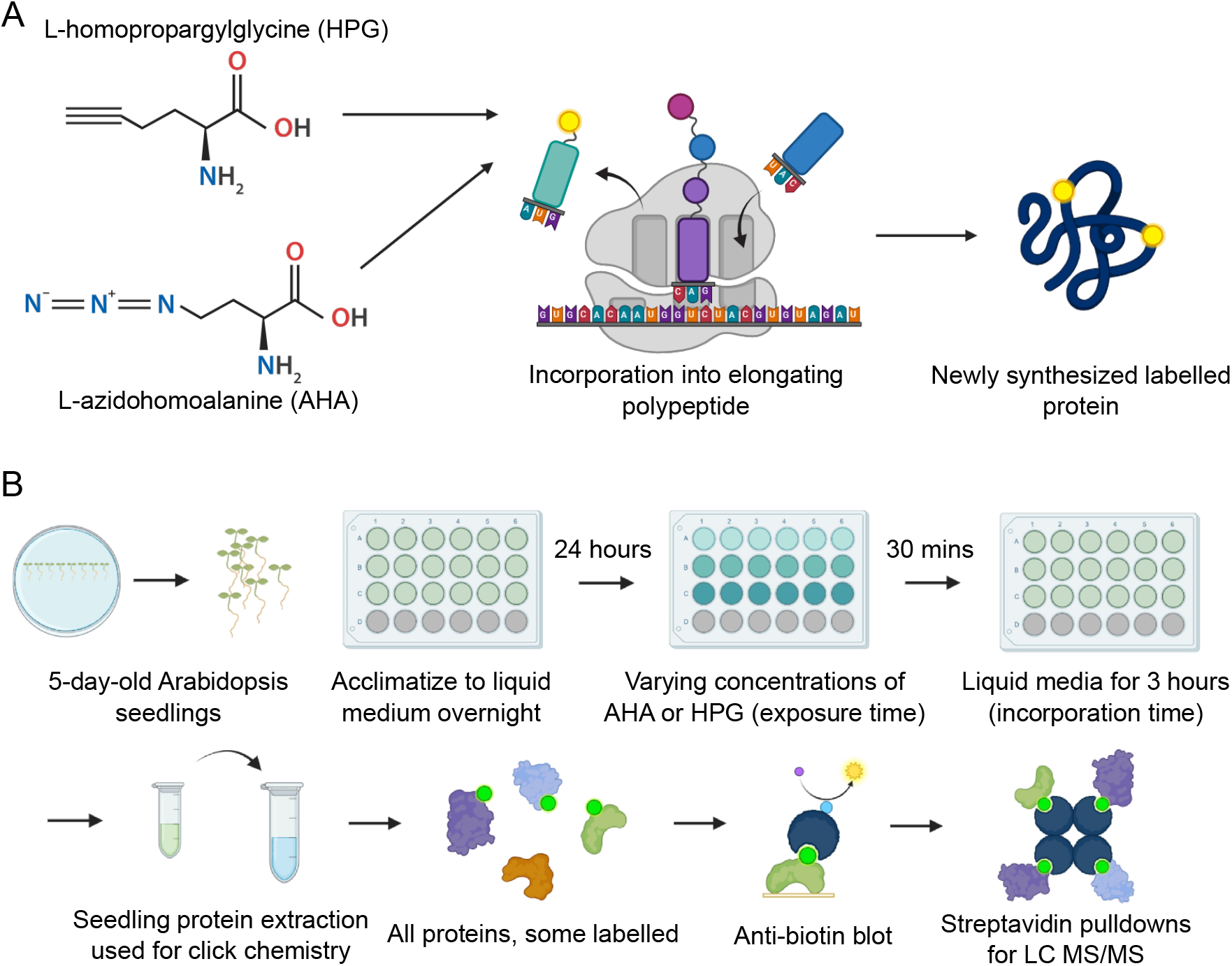
Visualization of BONCAT technology. A) Incorporation mechanism of AHA and HPG into the nascent proteome. B) Workflow for click labelling. Whole seedlings are acclimatized to liquid 0.5X MS media for 24 hours before treatment with AHA or HPG. Acute exposure time in the modified protocol is limited to 30 minutes, followed by 3 hours of incorporation time. Protein is extracted from seedlings for click chemistry in vitro, followed by visualization via anti-biotin blot to determine qualitative labelling before streptavidin enrichment for LC-MS/MS.

In the model plant *Arabidopsis thaliana* (Arabidopsis), BONCAT was first implemented in 2017. Here, AHA and HPG were applied to Arabidopsis seedlings through the flooding of agar plates with 1 mM AHA (Glenn et al., 2017). This yielded thousands of enriched proteins after only a few hours of exposure. However, subsequent endeavours found that Arabidopsis reacts metabolically to NCAAs similar to excess methionine, stunting plant growth and/or impacting plant health during prolonged exposure (Tivendale et al., 2021). Since then, only a handful of studies have utilized BONCAT in plants, with protocol updates generally focused on seedling exposure time to high concentrations of NCAAs (Li et al., 2023; Linster et al., 2022; Lyu et al., 2023; Xie et al., 2023; Yu et al., 2020). Together, these studies indicate that optimization of BONCAT in plants remains incomplete, as a wider variety of experimental parameters need to be tested to strike a balance between effective NCAA labelling concentrations and minimal disturbance plant health. Additionally, protocol simplification efforts remain unexplored, including: a) using the same biotin-conjugated clicked extract for both anti-biotin blots and enrichment for LC-MS/MS analysis to minimize variability and b) exploring the exclusivity of the azide-alkyne cycloaddition, including eliminating the need for repeated precipitation or sample processing prior to remove metabolites prior to “clicking” (Ma et al., 2018).

Here, we present an adapted protocol for BONCAT labelling of Arabidopsis seedlings in liquid MS medium that allows for more reliable seedling exposure to, and removal of, AHA and HPG (Figure 1B). Specifically, we demonstrate that the titration of AHA and HPG as well as optimization of labelling time is an effective approach to balance labelling while minimizing exposure to NCAAs. Lastly, we perform extensive seedling viability assays to ensure minimal disruption of plant growth using the modified protocol, and evaluate the newly synthesized proteome using the modified protocol under high and low acute salt and mannitol stress via LC-MS/MS.

## RESULTS AND DISCUSSION

### AHA and HPG label Arabidopsis seedlings at 50 µM or less using short application pulses

To determine concentrations of AHA and HPG suitable for application pulses, 5 d-old seedlings were acclimated to liquid medium overnight before being exposed to 0, 10, 25, 50, 100, or 250 µM AHA or HPG for 30 min (see Materials and Methods). Washed seedlings were then allowed to incorporate NCAAs into the nascent proteome for an additional 3 h before whole plants were flash frozen and extracted. Copper-catalyzed click chemistry was then performed *in vitro* to conjugate NCAA tags to biotin, and total proteome biotinylation was detected via immunoblot. Anti-biotin immunoblotting against untagged controls was selected as the preferred method for monitoring qualitative labelling efficiency, as whole proteome analysis may not detect AHA tags that have a mass shift similar to salt adducts of methionine (Liu et al., 2022). Here, AHA was found to have the best above-background labelling at all tested concentrations (Figure 2A). Based on these results, a concentration of 50 µM AHA or HPG was selected as an optimal concentration due to clearly visible labelling with minimal required reagent given the known metabolic stress of NCAAs (Tivendale et al., 2021).

**Figure 2.**
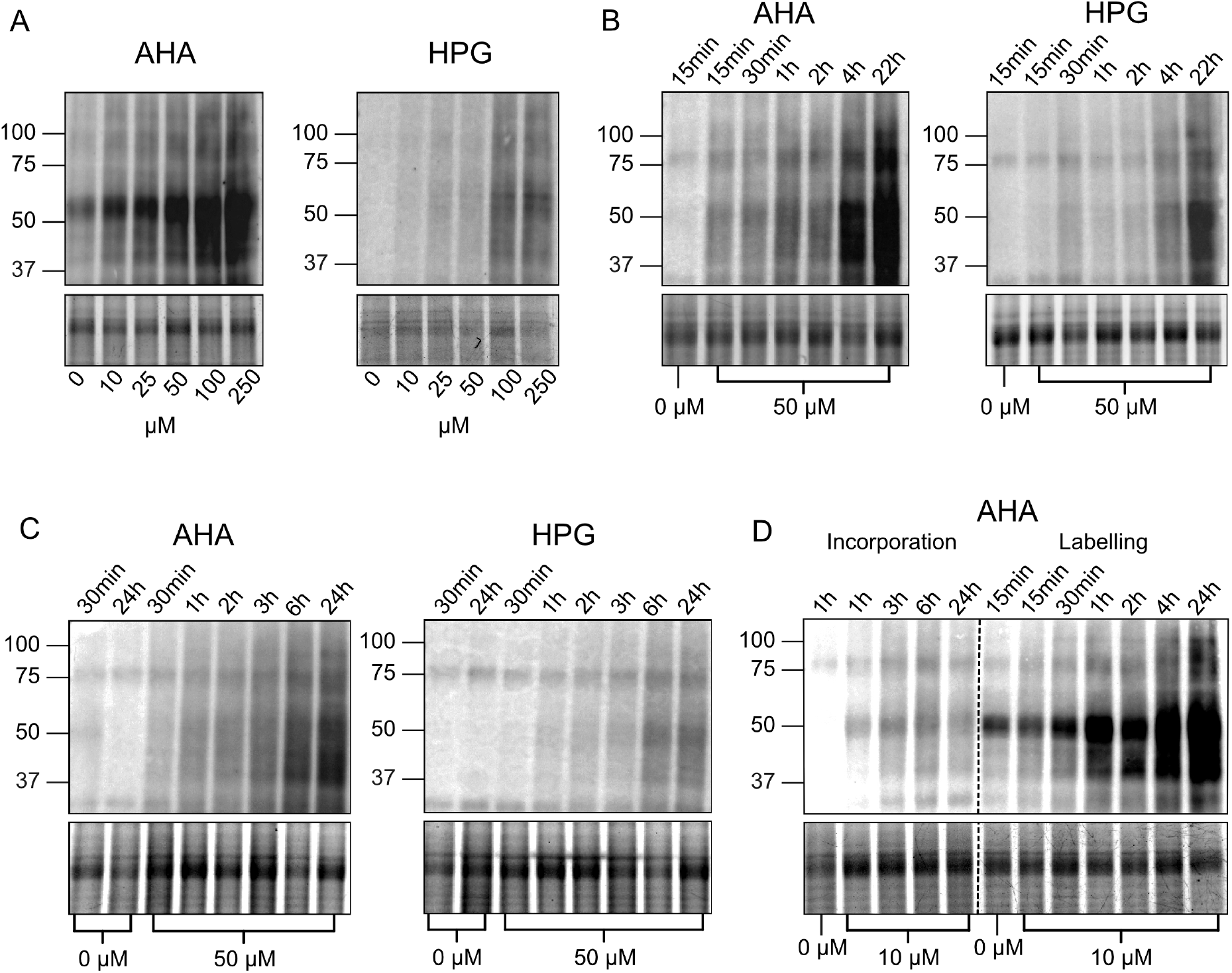
Streptavidin-detected anti-biotin blots of biotin-clicked Arabidopsis whole lysate samples. Beneath each blot is a cropped segment of a Coomassie Blue gel loading control. Each experiment was performed in triplicate, but single replicates are shown for simplicity. A) Titration of AHA and HPG concentrations with a 30-minute acute exposure and 3-hour incorporation time. B) 50μM AHA and HPG tested with exposure times varying from 15 minutes to 22 hours, all followed by a 3-hour incorporation. C) 50μM AHA and HPG tested with incorporation times varying from 30 minutes to 24 hours, each preceded with a 30-minute exposure time. D) Low concentration AHA (10μM) test of varying incorporation times with a 30-minute exposure (left) or varying exposure times with a 3-hour incorporation (right).

We then performed time-course labelling to select ideal labelling periods. Labelling periods were divided into acute exposure time (NCAA present in medium) and incorporation time (NCAA removed from medium), with NCAA proteome incorporation assessed independently. This revealed that increased exposure time increases labelling over the course of 24 h at both 50 µM AHA and HPG, with above-background labelling visible within the first 15 min (Figure 2B). Similarly, increased incorporation time with a set initial exposure of 30 min appears to increase labelling for up to 6 h after removal of tags using an initial 50 µM of either NCAA, suggesting residual NCAA likely remains present in the plant’s tissues after wash and removal from the medium, with that the remaining reagent continuing to be incorporated for several hours (Figure 2C). From this data, a 30 min exposure followed by 3 h of NCAA incorporation were selected as optimal labelling times for 50 µM of either NCAA, with a shorter acute time to mitigate potential negative effects of exposure on the seedlings. Interestingly, concentrations of AHA as low as 10 µM also showed increases in labelling across longer exposure times, but labelling stagnated with prolonged incorporation (Figure 2D). Therefore, a NCAA concentration of 10 µM is likely too low for excess NCAA to accumulate in plant tissue prior to its removal, but may be more suitable for long-term exposure labelling, such as overnight incubations for pulse-chase assay.

### Short exposure times prevent problems associated with prolonged exposure to AHA

Viability assays of NCAA treated Arabidopsis seedlings was performed by assessing fold-change in root length as a proxy for plant growth, in order to measure the potential negative effects of AHA and HPG. Here, 5 d-old Arabidopsis seedlings were transplanted onto 0.5x MS agar plates containing AHA or HPG and allowed to grow for multiple days. All tested concentrations from 10 to 250 µM AHA and HPG decreased plant root growth across a 3-day exposure, with HPG being significantly worse than AHA, even at comparably low concentrations (Figure 3A, Supplemental Figure 1A). However, with only 24 h of exposure, 10 µM AHA did not show a significant reduction in root growth. As this concentration shows above-background labelling within a few hours but does not continue to incorporate the NCAA after being washed (Figure 2D), 10 µM AHA may find an appropriate use in pulse-chase assays, where its capacity for 24-h acute exposure labelling could be used to label plants overnight for degradation experiments the following day.

**Figure 3.**
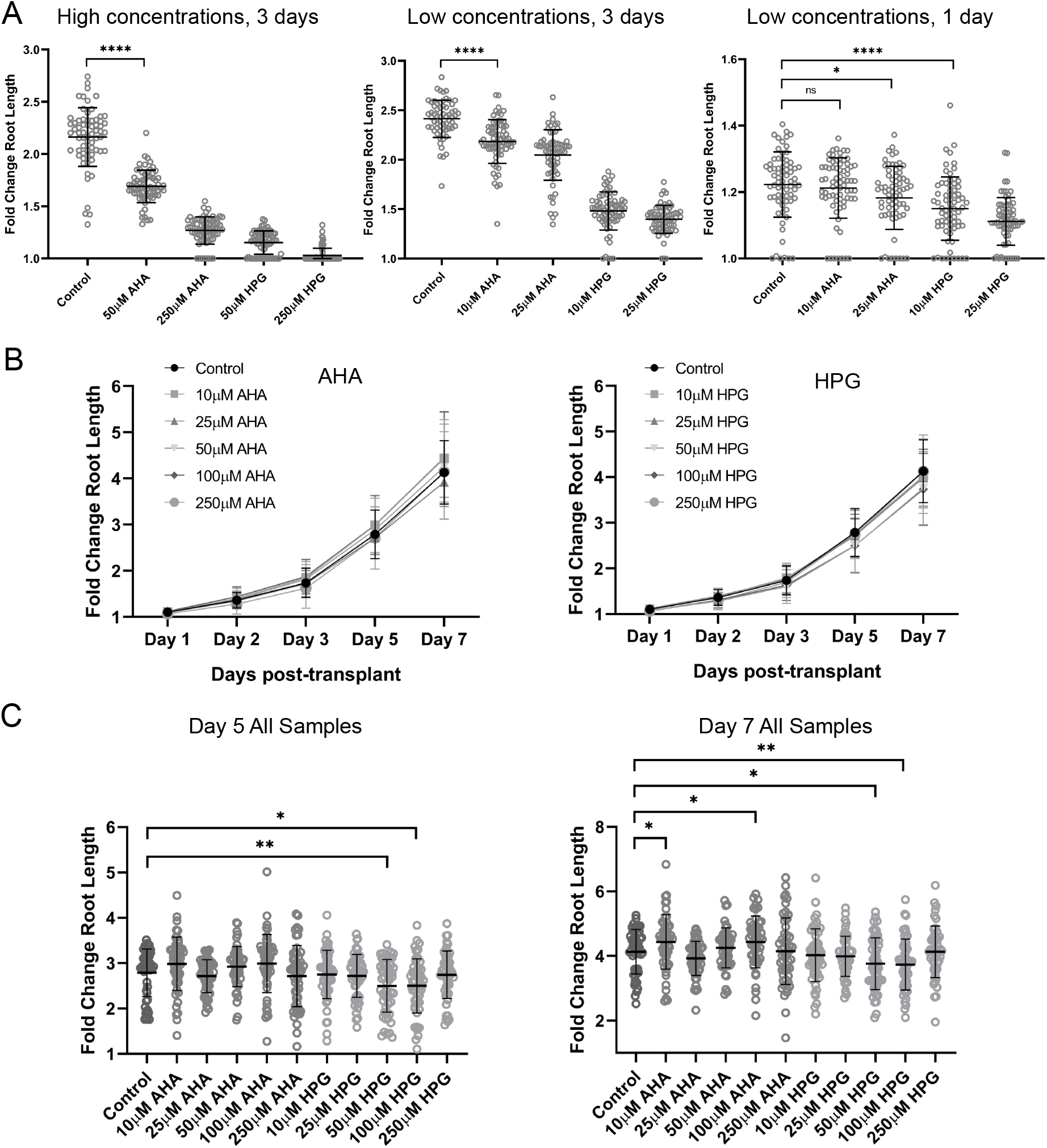
Viability assays of Arabidopsis seedlings exposed to methionine analogs. Average fold change in root length of seedlings with standard deviation error bars is plotted, and t-tests between experimental and control groups were performed with p-value significance < 0.05 shown. A) n = 72 5 d (days)-old seedlings per condition were transplanted to agar plates containing AHA or HPG for 1-3 d to simulate extensive exposure times. High concentrations after 3 days (left), low concentrations after 3 days (middle) and low concentrations after 24 hours (right) are shown. B) n = 54 seedlings per condition were acclimatized to liquid media for 24 h, then exposed for 30 min in AHA- or HPG-containing liquid media before being returned to solid medium and grown for 7 d. C) Snapshot of days 5 and 7 for all conditions from summarized data in 2B to show significant changes.

Differing approaches for *in vivo* AHA and HPG viability treatments have been deployed previously. For example, use of 1 mM AHA or HPG to acutely flood Arabidopsis seedlings growing on MS-agar plates revealed a negative impact on whole seedling growth and cell culture viability, in addition to perturbed metabolism (Tivendale et al., 2021). However, other eukaryotes demonstrate a more mixed or neutral response to NCAA exposure. For example, 2 mM AHA and 1 mM HPG have no negative viability effects on monkey and mouse-derived cells, while HeLa cells exhibit perturbation responses to AHA that seem to be mitigated by the presence of methionine in the growth medium (Bagert et al., 2014; Su Hui Teo et al., 2016; Wisse et al., 2017). On the molecular level, mouse-derived cell lines exhibit moderate stress signals (e.g. increased HEAT SHOCK PROTEIN 70 and 90 abundance) when treated with 50 µM AHA, but no significant increase in global protein ubiquitination and/or apoptosis/cell death associated with a reduction in viability (Morey et al., 2021).

In contrast, Tivendale et al. (2021) found AHA to be more toxic than HPG, especially in plant cell cultures. Bacteria have shown similar concentration-dependent sensitivity to NCAAs, but in a manner more consistent with this study. AHA appears to be tolerable to log-phase *E. coli* at higher concentrations than HPG, where HPG reduced bacterial growth at concentrations above 2.8 µM while AHA was tolerated up to the 200 µM range for prolonged exposure (Landor et al., 2023). This effect differs slightly between prototrophs and auxotrophs, where bacteria unable to synthesize essential compounds appear to be more negatively affected by HPG (Jecmen et al., 2023). Both AHA and HPG mildly affect bacterial metabolism as well, but are not degraded as the expected products of methionine (Steward et al., 2020).

It has been proposed that negative long-term effects of NCAAs on organism or cell growth may be irrelevant if incorporation into nascent proteins and immediate toxicity does not occur during the length of the experiment (Landor et al., 2023). To date, viability assays in plants have examined organism or cellular responses over prolonged exposure times, which is not always representative of the experimental conditions. Therefore, we repeated viability assays with a more representative exposure (short acute exposure in liquid 0.5x MS), followed by seedling washing and return to 0.5x MS - agar plates. All tested concentrations of AHA (up to 250 µM) did not reduce plant growth compared to controls after a 30-min AHA exposure (Figure 3B-C, Supplemental Figure 1B), suggesting that the long-term exposure growth defects are not a result of immediate toxicity of the NCAA. In contrast, 50 µM and 100 µM concentrations of HPG showed a slight reduction in growth compared to control under the same conditions (Figure 3C). By limiting exposure to 30 min and allowing incorporation to continue for < 6 h, viability concerns can be avoided without compromising in labelling efficiency, particularly in the use case of AHA. To strike a balance in labelling efficiency while minimising potential toxicity, the experimental settings for stress experimentation were set as 30 minutes of direct exposure to 50µM AHA, followed by 3 hours of incorporation time, and harvesting.

### AHA-labelled proteins under salt and osmotic stress

To benchmark the quantitative labelling efficiency of the optimized protocol under different experimental conditions, we applied two abiotic stress conditions during AHA labelling to compare conditions known to have overlapping, but not fully redundant, effects on the plant proteome (Rodriguez Gallo et al., 2023). The experiment was performed with either ‘high stress’ (150 mM salt or 300 mM mannitol) or ‘low stress’ (50 mM salt or 100 mM mannitol) conditions, as the effects of applied stressors may be intensified by liquid submersion. All stressors were applied for the entire duration of exposure and incorporation treatment (3 hours). Here, we find 549 proteins enriched in at least one condition under high stress conditions, 797 under low stress conditions, with a greater percentage of overlapping proteins between groups under lower stress versus high stress (55.1% vs. 35.1%; Figure 4A). Hundreds of these proteins represent Class I hits, which we define as below detection in the MS2 spectra of negative control samples (Figure 4B). Further principal component analyses found AHA-treated samples cluster away from untreated (bead binding control) samples, but distance between clusters of stressed and unstressed samples vary with abiotic stress intensity (Figure 5A and B).

**Figure 4.**
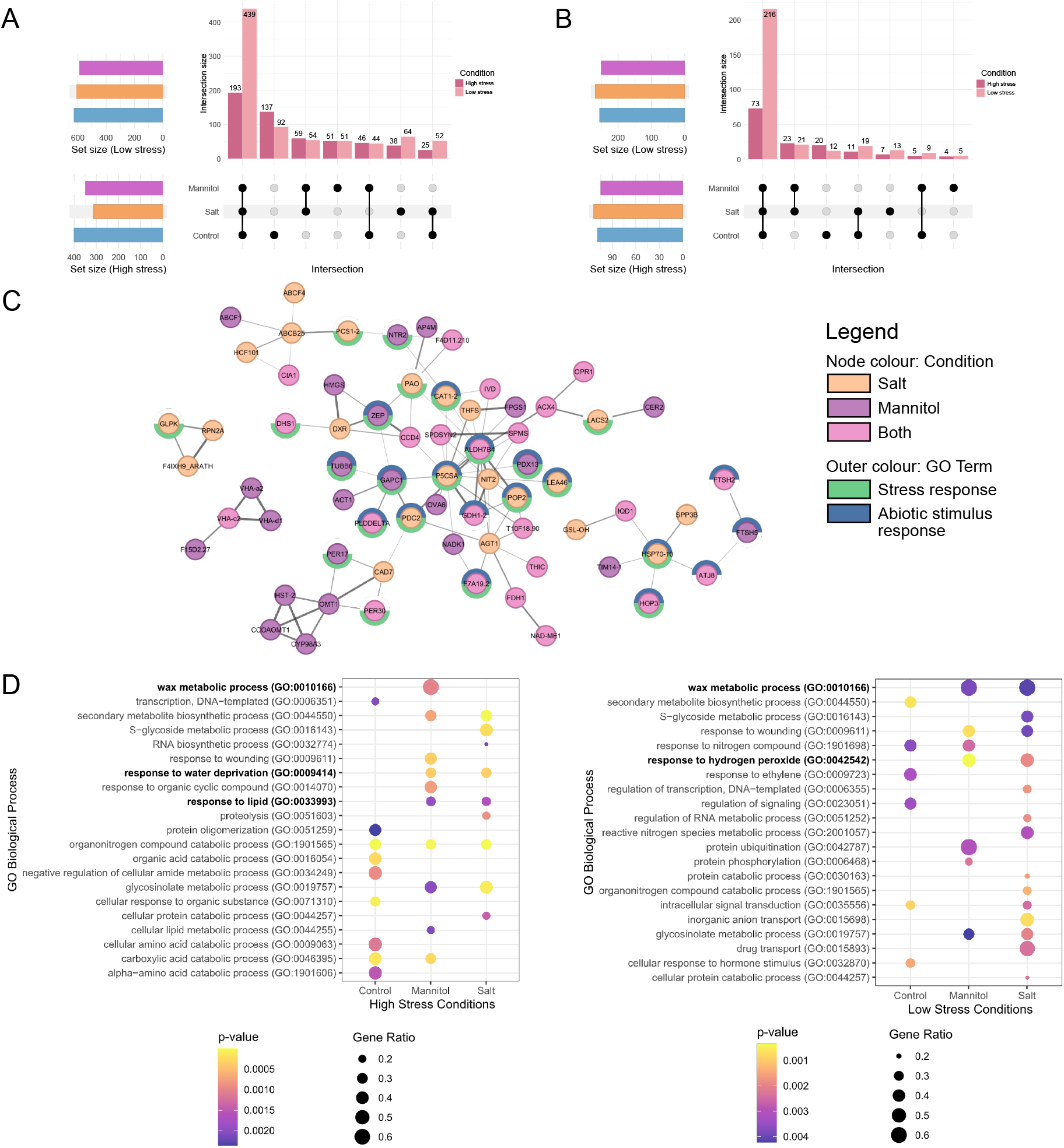
Mass spectrometry data for 50 μM AHA-labelled 3-hour salt or mannitol stressed Arabidopsis seedlings. A) Upset plot of enriched (>1.5 FC or below detection in negative control) protein groups and overlap between positive control (AHA treated, unstressed) and stress conditions in either high stress (150mM salt and 300mM mannitol, dark, n = 549) or low stress (50mM salt, 100mM mannitol, light, n = 797) conditions. B) Exclusive Class I hits from A that are absent in negative controls (n = 147 high stress and n = 300 low stress). C) STRING-DB association network of signficantly changing, enriched proteins from the low stress salt and/or mannitol conditions. Only nodes with > 2 edges are shown. Increasing line thickness is edge score ranging from 0.5-1. D) Gene ontology biological processes (Benjamini-Hochberg parent-child union) for high stress (left, p < 0.0025) and low stress (right, p < 0.005) conditions. All proteins enriched in one condition over negative controls were included. Highlighted in red are terms directly related to stress response.

**Figure 5.**
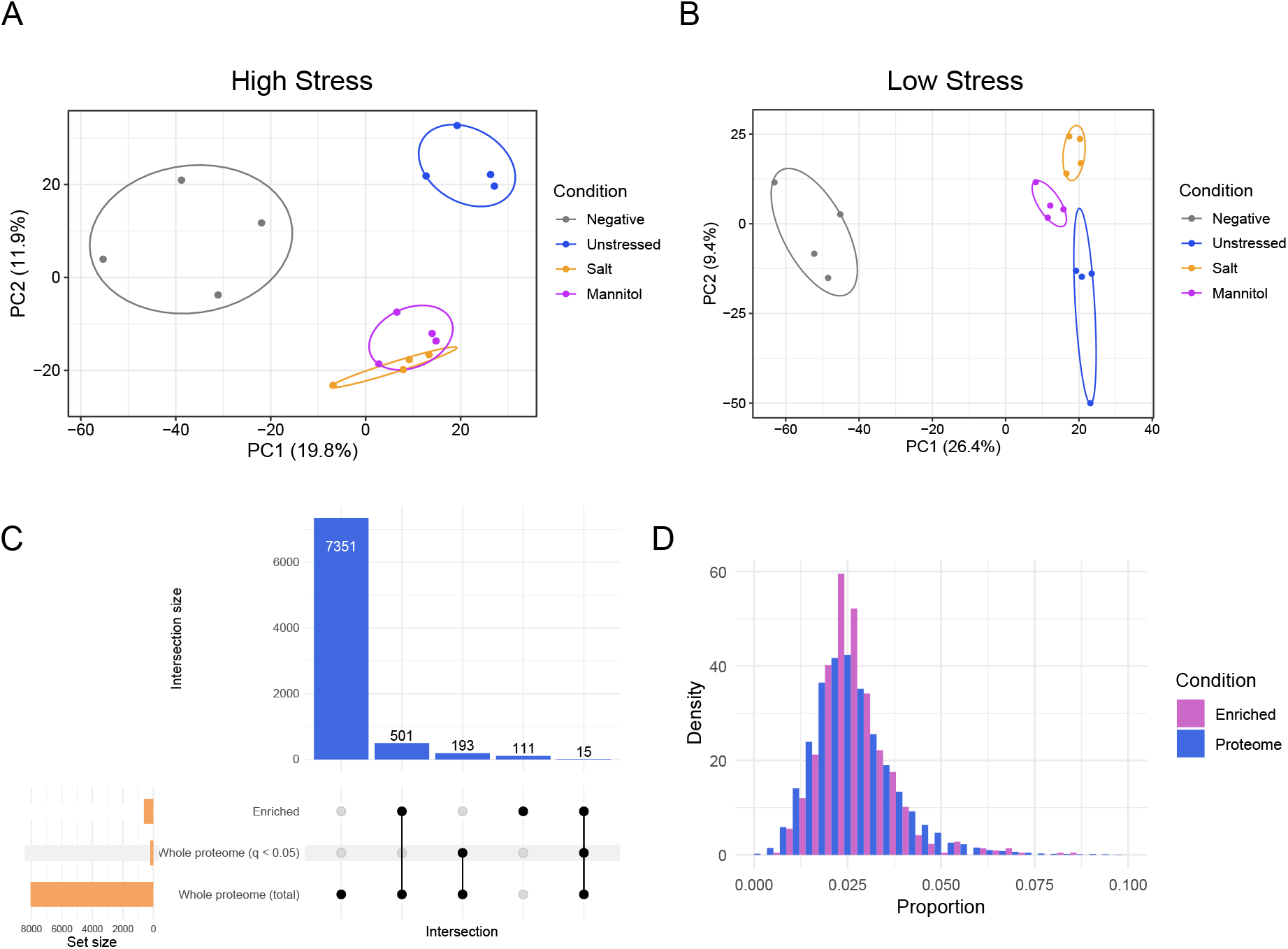
Proteomics data representation of salt and mannitol stressed AHA-treated seedlings. A) PCA plot of high stress (150mM salt - orange and 300mM mannitol - purple) 50µM AHA-treated seedlings, along with control treated (blue) and control untreated (grey) samples. Elipses are 68% confidence; n = 4. B) Same as A but with low stress samples (50mM salt and 100mM mannitol). C) Upset plot comparing all protein groups between the AHA-treated whole proteome (8060 proteins), significantly changing proteins in the whole proteome of AHA-treated samples vs. untreated whole proteome (208, > 0.58 Log_2_FC and *q-value < 0*.*05*), and significantly changing click-enriched proteins from AHA-treated seedlings vs. untreated streptavidin bead binding control (627, > 0.58 Log_2_FC and *q-value < 0*.*05*); n = 4. D) Comparison of relative methionine content (number of methionine residues over total length of protein) between the entire Arabidopsis proteome from Araport 11 (Proteome) and the 627 significant AHA-tagged proteins enriched (Enriched). Density represents a factor of population size to give the curves equal area.

Additionally, using positive AHA-labeled control samples, we can disentangle stress specific responses. Network analysis of dynamically changing salt and mannitol responsive proteins, resolved multiple protein networks related to abiotic stimulus response (Figure 4C). Gene ontology (GO) enrichment analysis further validated enrichment of abiotic stress response biological processes under the stress conditions, including wax biosynthesis and water-related regulation (Figure 4D, Supplemental Figure 2, Supplemental Figure 3). Consistent with prior findings, responses to salt and mannitol at the protein level appear to differ from each other (Supplemental Figure 4; Rodriguez Gallo et al., 2023) Next, we compared our BONCAT enrichment data to the changing salt and osmotic proteome resolved by Rodriguez Gallo et al. (2023). While BONCAT enrichment resolves overall fewer significantly changing proteins, we find hundreds of proteins in our BONCAT enrichment data stress conditions that were also significantly changing in the stress data of Gallo and colleagues (2023), including known salt and osmotic responsive CALCIUM-DEPENDENT PROTEIN KINASE 6 (AtCDPK6; Xu et al., 2010) and osmotic stress responsive RNA-binding protein ALDEHYDE DEHYDROGENASE 7B4 (AtALDH7B4; Figure 4C, Supplemental Figure 5A, Marondedze et al., 2019). We also find more than 100 proteins that while significantly enriched in our data under stress conditions that do not exhibit a significant change in their whole proteome abundance, despite gene ontology showing their involvement in abiotic stress (Rodriguez Gallo et al., 2023, Supplemental Figure 5B, Supplemental Figure 6). This indicates that BONCAT provides abiotic stress induced protein production response information that cannot be achieved through conventional whole proteome analyses, offering a more nuanced understanding of abiotic stress response relative to whole proteome analysis alone.

### BONCAT isolates newly produced proteins and proteins eluding quantification by whole proteome analyses

An advantage of BONCAT is the ability to observe proteins that may be degraded and produced at similar rate, or proteins produced in low abundance, both of which would go unnoticed in conventional total proteome analysis. To examine this scenarios, we examined comparable AHA-labeled and untreated control whole seedling proteomes to determine if there is a detectable effect of the amino acid analog treatment at the molecular level and to compare the proteins found in AHA-labelled whole proteome versus BONCAT enriched samples. Of the more than 8000 proteins quantified in the whole proteome, only 208 were significantly different between AHA-treated and untreated control samples (∼2.5%; *q-value* < 0.05), with only 15 of those significant proteins detected in the AHA enriched sample set of 628 (∼2.3%; Figure 5C), indicating at low concentrations AHA elicits minimal molecular perturbations. Furthermore, of the 628 AHA-treated proteins enriched without added stress conditions, we quantified 501 to not be significantly changing in the whole proteome, indicating that AHA labelling is precisely detecting the production of new proteins. Additionally, 111 of the 628 BONCAT-enriched proteins are not detected in the whole proteome analysis, indicating click-based enrichment allows detection of newly synthesized proteins that are low in abundance. Lastly, we examined our click-enriched proteins for bias towards methionine rich proteins (Figure 5D). Here, no bias for methionine rich proteins relative to the Arabidopsis proteome was found. Collectively, these results further indicate that we are enriching for proteins that are produced in response to salt stress.

## SUMMARY

In this study, we established a simple and usable workflow for the implementation of BONCAT in plants. Although 50 µM AHA was selected as the optimal labelling concentration, the protocol is flexible and modifiable based on need, with modified protocols able to be tracked for efficiency at multiple stages. Our BONCAT implementation showed an ability clearly define stress and unstressed conditions in relatively short periods that mitigates potential negative plant responses. This unveils outstanding new opportunities for applications where enrichment of a subsect of the proteome is required for sufficient depth of analysis. Lastly, the potential for NCAA-tag incorporation at concentrations substantially below the literature standard has positive new implications for the use of BONCAT in plants moving forward.

## MATERIALS AND METHODS

### Plant growth conditions

*Arabidopsis thaliana* seeds (Col-0) were liquid sterilized by immersion in 70% (v/v) ethanol for 2 min followed by 30% (v/v) bleach for 10 min. Sterilized seeds were vernalized in H_2_O in the dark at 4°C for 3 d. Germination plates were prepared using half-strength MS basal salts (Caisson Laboratories Inc. Murashige & Skoog MSP01), pH adjusted to 5.8 with aqueous KOH, with 0.08% (w/v) agar (Caisson laboratories Inc. Phytoblend PTP01).

For click chemistry, vernalized seeds were stratified on plates and allowed to grow with plates positioned vertically in 24 h light at 22°C for 5 days. Approximately 100 mg of 5 d-old seedlings were immersed in autoclaved half-strength MS media in 24-well plates and placed under constant light for 16 h to acclimatize. MS media was removed and replaced with 50 µM filter sterilized L-azidohomoalanine or L-homopropargylglycine (Vector Labs, CCT-1066 and CCT-1076) from 0.1 M stocks dissolved in H_2_O and diluted in 0.5x MS media. Seedlings were then incubated for 30 min of exposure time in constant light before being washed once in 0.5x MS and incubated in constant light in 0.5x MS for 3 hours of “incorporation” time. Stress experiments included either salt (150mM in ‘high stress’ and 50mM in ‘low stress’ conditions) or mannitol (300mM in ‘high stress’ and 100mM in ‘low stress’ conditions) in the 0.5X MS liquid media during both exposure and incorporation phases. Labelled seedlings were removed from media, rinsed in 0.5x MS, and quickly touch dried before being flash frozen in liquid N_2_ and stored at −80°C until subsequent steps.

To perform root length assays, seeds were sown individually on 0.5x MS agar plates positioned vertically in dark boxes with seeds at the light interface to emulate roots growing in the dark and shoots growing in the light. Seedling growth conditions consisted of a 12 h photoperiod, 22°C and 100 µmol/m^2^s light. After 5 d for the long-term reagent exposure, seedlings were transplanted along the same light/dark interface to 0.5x MS plates containing 0, 10, 25, 50, or 250 µM AHA or HPG. For the short-term exposure, seedlings were placed into liquid 0.5x MS overnight to equilibrate, then immersed for 30 min in 0.5x liquid MS containing 0, 10, 25, 50, or 250 µM AHA, before being washed in 0.5x MS and re-plated on to vertical plates. Root length was recorded for 7 d using digital calipers, and fold-change in individual seedling root length relative to day 0 (transplanting day) was plotted. Recording of root length stopped after day 3 for the long-term exposure due to statistically significant differences between all concentrations relative to controls.

### Optimization of plant labelling

To determine optimal concentrations of label, variable concentrations of 10, 25, 50, 100, or 250 µM of AHA or HPG were applied as above. Optimal concentrations for labelling were chosen based on a combination of intensity of labelled protein present on streptavidin immunoblots and minimal observed growth retardation on 0.5x MS agar containing the same concentration of label. For time-course experimentation, acute exposure times were varied between 15 min, 30 min, or 1, 2, 4, or 22 h. Incorporation times were varied between 30 minutes or 1, 2, 3, 6, or 24 h. Optimal incubation times were chosen based on minimal required times to achieve effective labelling at the desired concentration as visualized via immunoblot, and verified to cause insignificant or no growth inhibition to the seedlings based on viability studies.

### Protein extraction and quantification

Flash frozen tissue was ground using Geno/Grinder (SPEX SamplePrep) for 30 s at 1200 rpm with 3mm glass beads and extracted in either 50mM HEPES (pH 7.5) with 1% (w/v) SDS or 50 mM Tris (pH 8.0) with 1% (w/v) SDS, at a ratio of 1:2 w/v ground tissue to extraction buffer. Following optimization, HEPES was selected as the extraction buffer for all samples due to better click labelling observed. Samples were heated at 95°C for 10 min and 1000 rpm on Eppendorf ThermoMixer F2.0, cooled, and centrifuged at 16,000 xg for 10 min at room temperature to remove tissue particulates. Protein amounts were quantified using Bradford assay with samples diluted to 0.2% (w/v) SDS against BSA controls.

### Click chemistry reaction

A total of 250 µg of protein from each sample was diluted in extraction buffer with 1% (w/v) or no SDS to a volume of 346 µL ensuring a final concentration of 0.5% (w/v) SDS to maintain protein solubility. To each reaction, the following was added: 8 µL of 50 mM tris(2-carboxyethyl)phosphine (TCEP)-HCl (Sigma Cat. No: 51805-45-9; diluted from 1 M stock in H_2_O same day of use), 30 µL of 1.7 mM tris[(1-benzyl-1H-1,2,3-triazol-4-yl)methyl]amine (TBTA) (Vector Labs Cat. No: CCT-1061; in 20% (v/v) dimethyl sulfoxide (DMSO) and 80% (v/v) tert-butanol, stored at room temperature as in Ma et al., 2018Ma et al., 2018), 8 µL of CuSO4 (Sigma Cat. No: 451657-10G; in H_2_O, stored at −20°C), and 8 µL of 5mM biotin-PEG3-azide (Vector Labs Cat. No: CCT-AZ104; for HPG samples, in DMSO) or biotin-PEG4-alkyne (Vector Labs Cat. No: CCT-TA105; for AHA samples, in DMSO). Click reactions were allowed to proceed for 1 h at room temperature, shaking at 700 rpm and protected from light. Reactions were then quenched via chelation with 20 µL of 0.5 M ethylenediaminetetraacetic acid (EDTA) (pH 8.0, stored at 4°C). Labelling validation was determined via streptavidin immunoblot using 6 µg of protein run on 10% SDS-PAGE gels transferred to 0.45 µM polyvinylidene fluoride (PVDF) membrane. Blots were blocked in 5% fat-free milk in Tris-buffered saline (TBS) for 1 h, at which point HRP-conjugated streptavidin (Thermo Scientific Cat. No. N100) diluted 1:2000 in Tris-buffered saline with 0.1% Tween-20 (TBST) was added for an additional hour. Membranes were washed in TBST and visualized using Clarity Western ECL Substrate (Bio-Rad Cat No. 170-5060). Colloidal Coomassie Blue-stained gels were used as loading controls.

### Enrichment of clicked proteins

Excess biotin and small molecules were removed from clicked samples using Cytiva PD Spintrap G-25 columns (Cat. No: 28918004). Samples were bound to Cytiva streptavidin Sepharose beads (Cytiva Cat. No. 17511301); all depicted experiments proceeded using non-magnetic Sepharose beads. All beads were initially washed 3 times in 50 mM HEPES, pH 7.5, for 1 min while rotating at room temperature. For all steps, supernatant was removed by centrifugation for 1 min at 600 xg. Up to 200 µg of flow-through protein from the Spintrap columns was loaded onto equilibrated streptavidin beads and allowed to bind while rotating for 16 hours at 4°C. The supernatant was removed and retained at −20°C to assess capture efficiency. A total of 4 biological replicates were enriched in each experimental condition.

Capture efficiency was checked before digestion via streptavidin-detected dot blot comparison of clicked protein, Spintrap column flow-through, and supernatant after overnight bead binding. After confirmation of a signal reduction of more than 70% in the supernatant after binding, beads were washed sequentially with the following solutions at 1mL each: 50 mM HEPES pH 7.5 (2x washes), 1 M potassium chloride, ice-cold 100 mM sodium carbonate (2x washes), 2 M urea in 50 mM Tris-HCl pH 8.0 (2x wash), and 50 mM triethylammonium bicarbonate (TEAB; Sigma Cat. No: 15715-58-9) in HPLC-grade water (3x washes). All washes were completed at room temperature with rotation for 3 min.

### Trypsin digestion and peptide processing

*Enriched Proteome -* Peptides were generated through on bead digestion using sequencing grade trypsin (V5113; Promega) diluted 1:100 in 50 mM TEAB overnight at 37°C; the supernatant was removed before the beads were washed with 50 mM TEAB and the supernatant pooled with previously extracted peptides. The extract was reduced with 10 mM DTT at 95°C for 5 min while shaking at 450 rpm, alkylated with 30 mM iodoacetamide (IA) at room temperature for 30 min, and the reaction quenched with 10 mM DTT for 10 minutes. Samples were then acidified to 0.5% (v/v) TFA to a pH of less than 2 and centrifuged at 16,000 xg for 1 min before the supernatant was removed and dried down. Peptides were re-suspended in 3% (v/v) acetonitrile / 0.1% (v/v) formic acid for desalting via ZipTip (Sigma Cat. No. ZTC18S960) before being dried and stored at −80°C. Desalted peptides were the re-suspended in 3% (v/v) acetonitrile / 0.1% (v/v) formic acid for mass spectrometry analysis.

*Whole Proteome -* 100 µg of Arabidopsis seedling protein extracts were generated as previously described without deviation (Rodriguez Gallo et al. 2023). Samples were then digested and processed as described above.

### LC MS/MS and Data Analysis

Peptides were analyzed using a FAIMSpro mounted Fusion Lumos Orbitrap mass spectrometer (Thermo Fisher Scientific) in data independent acquisition (DIA) mode. Peptides were injected using Easy-nLC 1200 system (LC140; Thermo Fisher Scientific) and an Acclaim PepMap 100 C18 trap column (Cat# 164750; Thermo Fisher Scientific) followed by a 50 cm Easy-Spray PepMap C18 analytical column (ES903; Thermo Fisher Scientific) warmed to 50 °C. Peptides were eluted at 0.3 μl/min using a segmented solvent B gradient of 0.1% (v/v) formic acid in 80% (v/v) acetonitrile from 4 to 41% solvent B (0–60min). The FAIMSpro was used with a fixed gas flow of 3.5 L/min with a 3-CV setting of −30, −50, −70. A positive ion spray voltage of 2.3 kV was used with an ion transfer tube temperature of 300 °C and an RF lens setting of 40%. Direct DIA acquisition was performed as previously described (Mehta et al., 2022). Full scan MS^1^ spectra (350–1400 *m*/*z*) were acquired with a resolution of 120,000 at 200 *m*/*z* with a normalized automatic gain control (AGC) target of 100% and IT set to automatic. Fragment spectra were acquired at resolution of 30,000 across 28 38.5 *m*/*z* windows overlapping by 1 *m*/*z* using a dynamic maximum injection time and an AGC target value of 2000%, with a minimum number of desired points across each peak set to 6. Higher-energy collisional dissociation fragmented was performed using a fixed 27% fragmentation energy. Downstream data analysis of dDIA acquisitions was performed using Spectronaut ver. 19 (Biognosys AG) with default settings as previously described (Mehta et al., 2022), with all searches made against a custom-made decoy (reversed) version of the Arabidopsis protein database from Araport 11 (27,533 protein encoding genes; ver. 2022-09-14). Briefly, search parameters included the following: trypsin digest permitting two missed cleavages, fixed modifications (carbamidomethyl (C)), variable modifications (oxidation (M)) and a peptide spectrum match, peptide and protein false discovery threshold of 0.01.

### Bioinformatic Data Analysis

Significantly changing proteins (Log_2_FC of MS2 spectra > 0.58 over negative control) were pooled with Class I protein hits (absent in MS2 of negative controls) to form the foreground with a background of all identified proteins (Supplemental Table 1, Supplemental Table 2). Comparison to whole proteome analysis was done to both AHA-treated unstressed whole proteomes that followed the same standards for analysis (Supplemental Table 3) and previous work on salt and mannitol stressed whole proteomes by Rodriguez Gallo et al. (2023; Supplemental Table 4). Gene Ontology (GO) enrichments were performed using the Ontologizer (Bauer et al., 2008; http://ontologizer.de/, Supplemental Table 5, Supplemental Table 6) using a Benjamini-Hochberg parent-child union with an analysis cutoff of p > 0.01, and visualized in R 4.5.0. Proteins below differential abundance in both positive and negative controls was used for network analysis of stress-exclusive proteins via the StringApp plugin for Cytoscape v. 3.10.3, with an edge score cut-off of 0.5 (Doncheva et al., 2019, https://cytoscape.org/). STRING Enrichment using *StringApp* was used to assign gene ontologies. Methionine content analysis was done using in-house Python scripts to analysis methionine content and length of the Arabidopsis protein database from Araport 11 (https://phytozome-next.jgi.doe.gov/ ; Supplemental Table 7). PCA analysis was performed using *SRplot* (Tang et al., 2023), with other visualizations produced using a combination of *ggplot2*, Biorender (https://www.biorender.com/), GraphPad Prism v. 8.4.3 (https://www.graphpad.com/), and Affinity Designer v. 1.10.5 (https://affinity.serif.com/en-us/designer/).

## ACKNOWLEDGEMENTS

The authors would like to thank the Canadian Foundation for Innovation (CFI) and Natural Sciences and Engineering Council of Canada (NSERC) for funding this work. We would also like to thank Jack Moore and the Alberta Proteomics and Mass Spectrometry Facility (APM) for mass spectrometry assistance and upkeep.

## AUTHOR CONTRIBUTIONS

*Nicholas Hassan*: Investigation, Conceptualization, Methodology, Data curation, Formal analysis, Visualization, Writing—original draft. *Shelly Braun*: Writing—review & editing, Conceptualization. *Mohana Talasila*: Formal analysis, Data curation, Visualization. *Curtis Kennedy*: Data curation, Visualization. *Luke Yaremko*: Formal analysis, Data curation, Visualization. *Richard Fahlman*: Supervision, Writing—review & editing, Material Resources. *R. Glen Uhrig*: Supervision, Conceptualization, Writing—review & editing, Funding acquisition.

## DATA AVAILABILITY

Raw data have been deposited to the ProteomeExchange Consortium (http://proteomecentral.proteomexchange.org) *via* the PRoteomics IDEntification Database (PRIDE; https://www.ebi.ac.uk/pride/) partner repository with the data set identifiers PXD064210

## SUPPLEMENTAL FIGURES

**Supplemental Figure 1. Representative images of seedling samples during AHA and HPG viability assays**.

**Supplemental Figure 2. Enlarged gene ontology of high stress set AHA-treated seedlings**.

**Supplemental Figure 3. Enlarged gene ontology of low stress set AHA-treated seedlings**.

**Supplemental Figure 4. Heatmap representation of shared proteins between salt and mannitol stressed AHA-treated seedlings**.

**Supplemental Figure 5. Comparison of salt and mannitol stress data between previous whole proteome analysis and AHA-tagged enrichments**.

**Supplemental Figure 6. Gene ontology of salt and mannitol stress data specific to AHA-tagged enrichments compared to previous whole proteome analysis**.

## SUPPLEMENTAL TABLES

**Supplemental Table 1. Enriched AHA-treated low and high salt and mannitol stress proteomics data**.

**Supplemental Table 2. Differentially expressed proteins above significance threshold divided by group between positive control, salt stressed, and mannitol stressed seedlings in both high and low stress**.

**Supplemental Table 3. Whole proteome data for AHA-treated and untreated proteomes**.

**Supplemental Table 4. Comparison of significantly changing proteins in AHA-treated stress experiments with significantly changing proteins in stress experiments by Rodriguez Gallo et al. (2023)**

**Supplemental Table 5. Ontologizer GO output data for AHA-treated low and high salt and mannitol stressed proteins**.

**Supplemental Table 6. Ontologizer GO output data for AHA-treated stressed proteins that are absent from previous whole proteome stress analysis**.

**Supplemental Table 7. Methionine proportion analysis raw output data between AHA-enriched proteins and the entire Arabidopsis proteome**.

