## Supplemental Figure 1 for "Optimization of Bio-Orthogonal Non-Canonical Amino acid Tagging (BONCAT) for effective low-disruption labelling of Arabidopsis proteins *in vivo*"

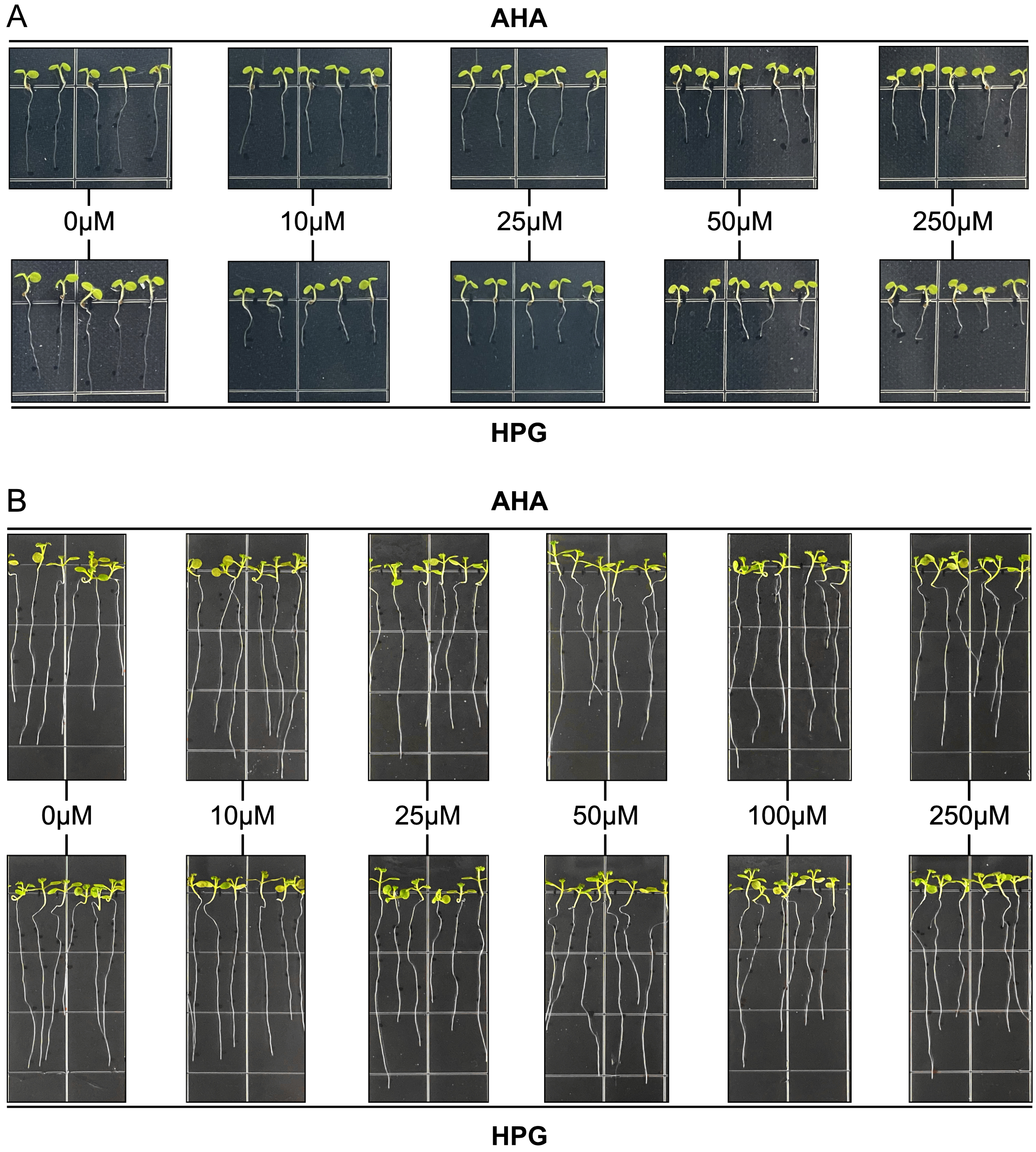


**Supplemental Figure 1. Representative images of seedling samples during AHA and HPG viability assays.** All seedlings were grown vertically on 0.5X MS agar for 5d before transplanting. Controls are dd$H_2O$. Images were taken at the end of each growth assay. A) Root length images at 3d post-transplant of seedlings with chronic exposure to AHA or HPG through 0.5X MS agar containing NCAAs. Full data is represented in Figure 3A. B) Root length images at 7d post-transplant of seedlings with brief (30min) exposure to AHA or HPG followed by wash and re-plating on 0.5X MS agar to verify incorporation phase viability. Full data is represented in Figure 3B and C.
