## Supplemental Figure 3 for "Optimization of Bio-Orthogonal Non-Canonical Amino acid Tagging (BONCAT) for effective low-disruption labelling of Arabidopsis proteins *in vivo*"

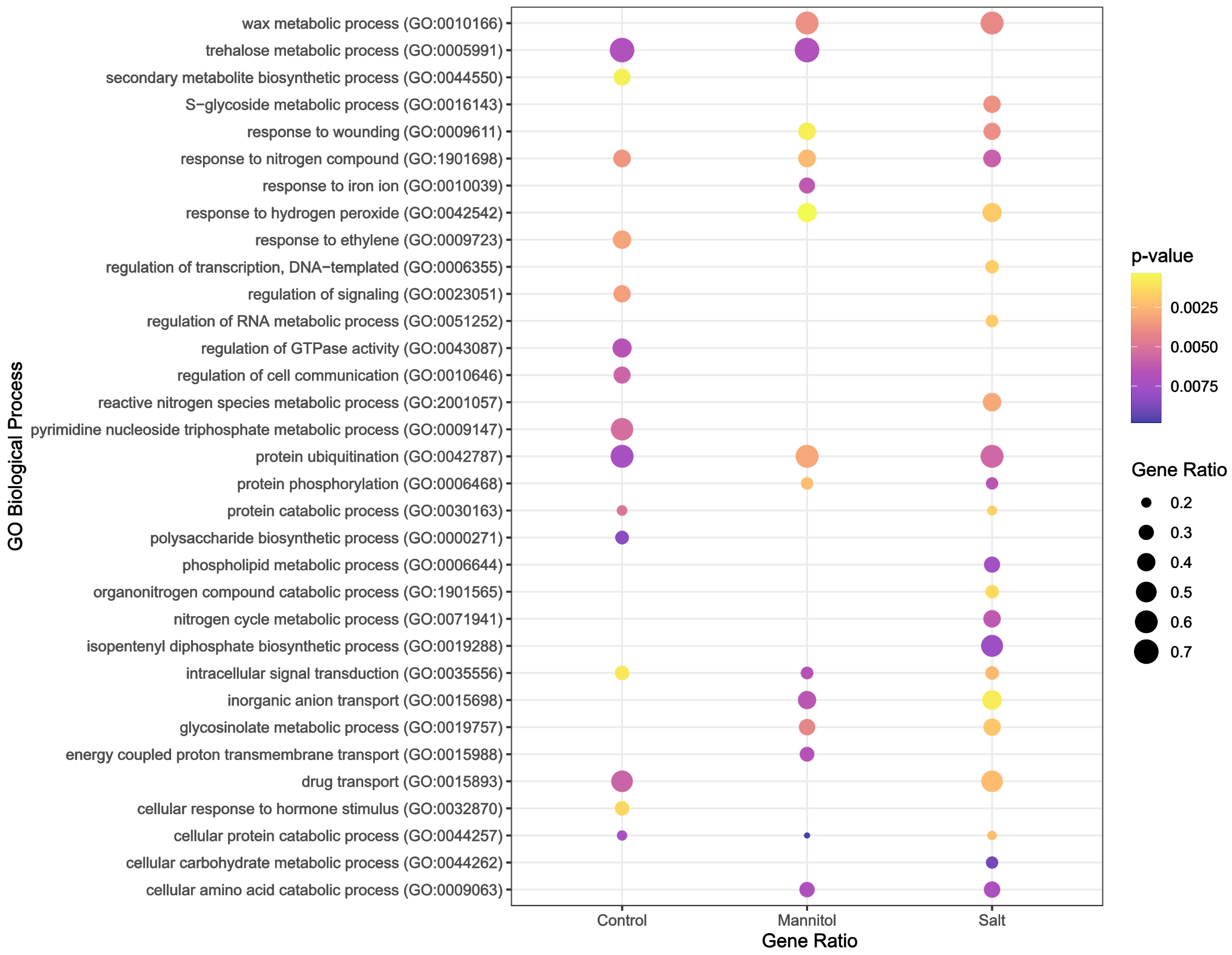


**Supplemental Figure 3. Enlarged gene ontology of low stress set AHA-treated seedlings.** GO terms with p-value < 0.01 (Benjamini-Hochberg parent-child union) for biological processes of proteins with $Log_2$ fold change > 0.58 over unlabelled controls in 50μM AHA-treated unstressed control, 50mM salt stressed, or 100mM mannitol stressed seedlings are represented.
