## Supplemental Figure 4 for "Optimization of Bio-Orthogonal Non-Canonical Amino acid Tagging (BONCAT) for effective low-disruption labelling of Arabidopsis proteins *in vivo*"

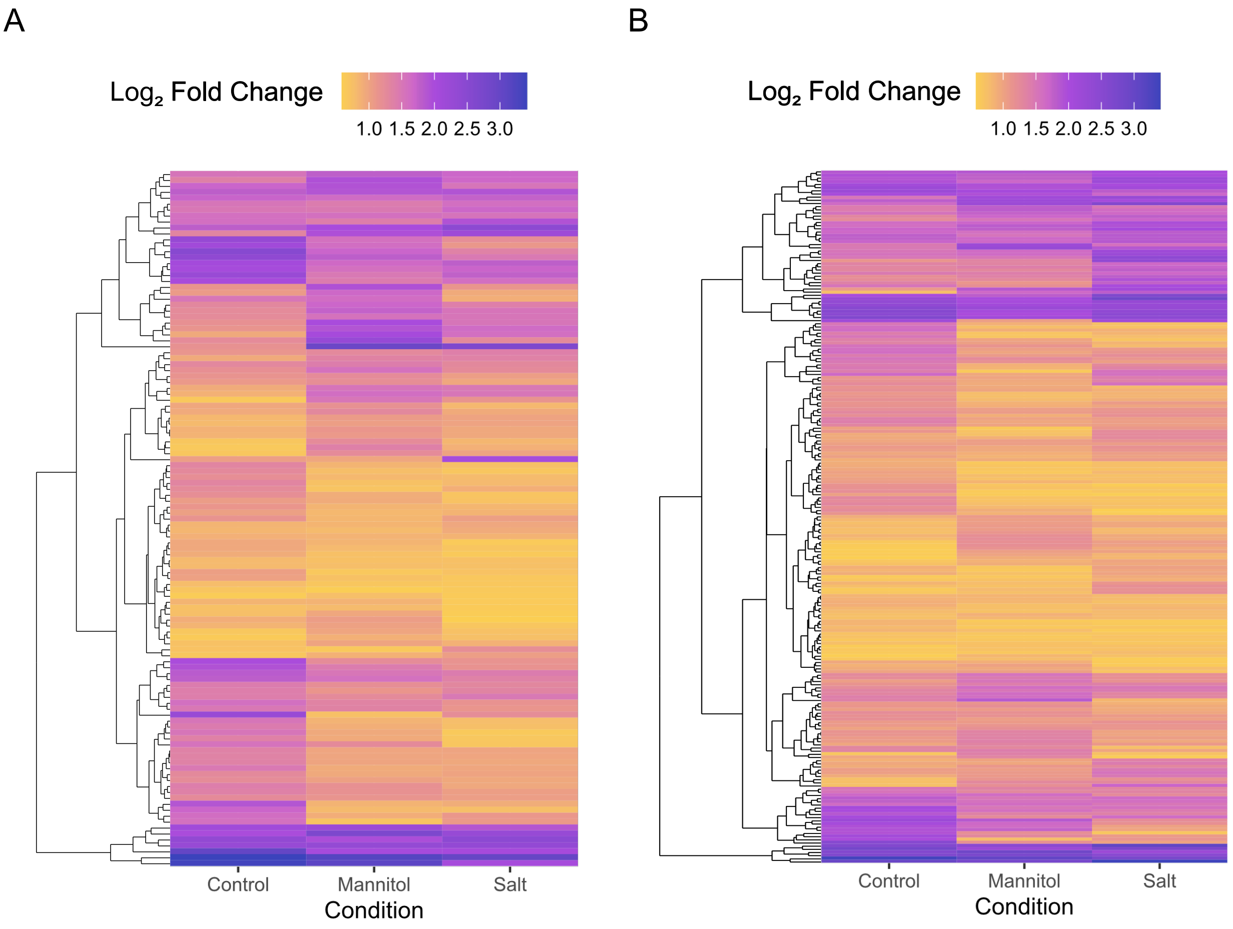


**Supplemental Figure 4. Heatmap representation of shared proteins between salt and mannitol stressed AHA-treated seedlings**. A) Euclidian distance-clustered heatmap of differentially expressed proteins that were significantly enriched (>0.58 Log$_2$ fold change) in all 3 AHA-treated conditions over untreated control in the high stress conditions, without Class I hits that were absent in the negative control (n = 117). B) Same as A, but for only terms enriched in all 3 groups in low stress conditions (n = 219).
