## Supplemental Figure 5 for "Optimization of Bio-Orthogonal Non-Canonical Amino acid Tagging (BONCAT) for effective low-disruption labelling of Arabidopsis proteins *in vivo*"

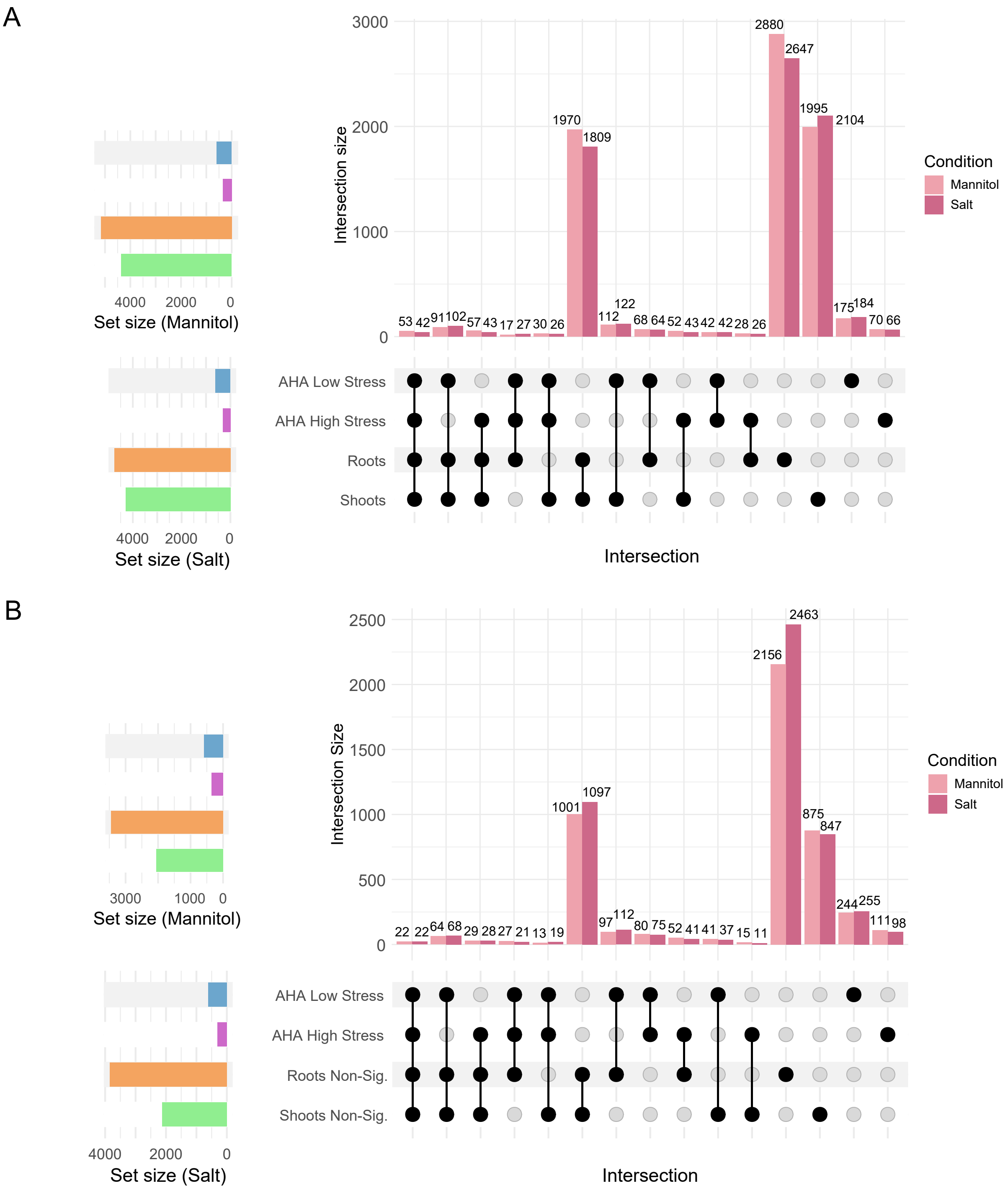


**Supplemental Figure 5. Comparison of salt and mannitol stress data between previous whole proteome analysis and AHA-tagged enrichments.** Data for whole proteome analysis was obtained by Rodriguez Gallo et al. (2023). **A)** Comparison of significantly changing root or shoot proteins under salt and mannitol stress (Rodriguez Gallo et al 2023; > 0.58 Log2FC and q-value < 0.05, n = 5) to significantly changing click-enriched proteins from AHA-treated seedlings during salt or mannitol stress (> 0.58 Log2FC and q-value < 0.05, n = 4). **B)** Upset plot of non-significant (q > 0.05) proteins from roots or shoots detected by Gallo et al. (n = 5) compared with significantly changing click-enriched proteins from AHA-treated seedlings during salt or mannitol stress (> 0.58 Log2FC and q-value < 0.05, n = 4).
