## Supplemental Figure 6 for "Optimization of Bio-Orthogonal Non-Canonical Amino acid Tagging (BONCAT) for effective low-disruption labelling of Arabidopsis proteins *in vivo*"

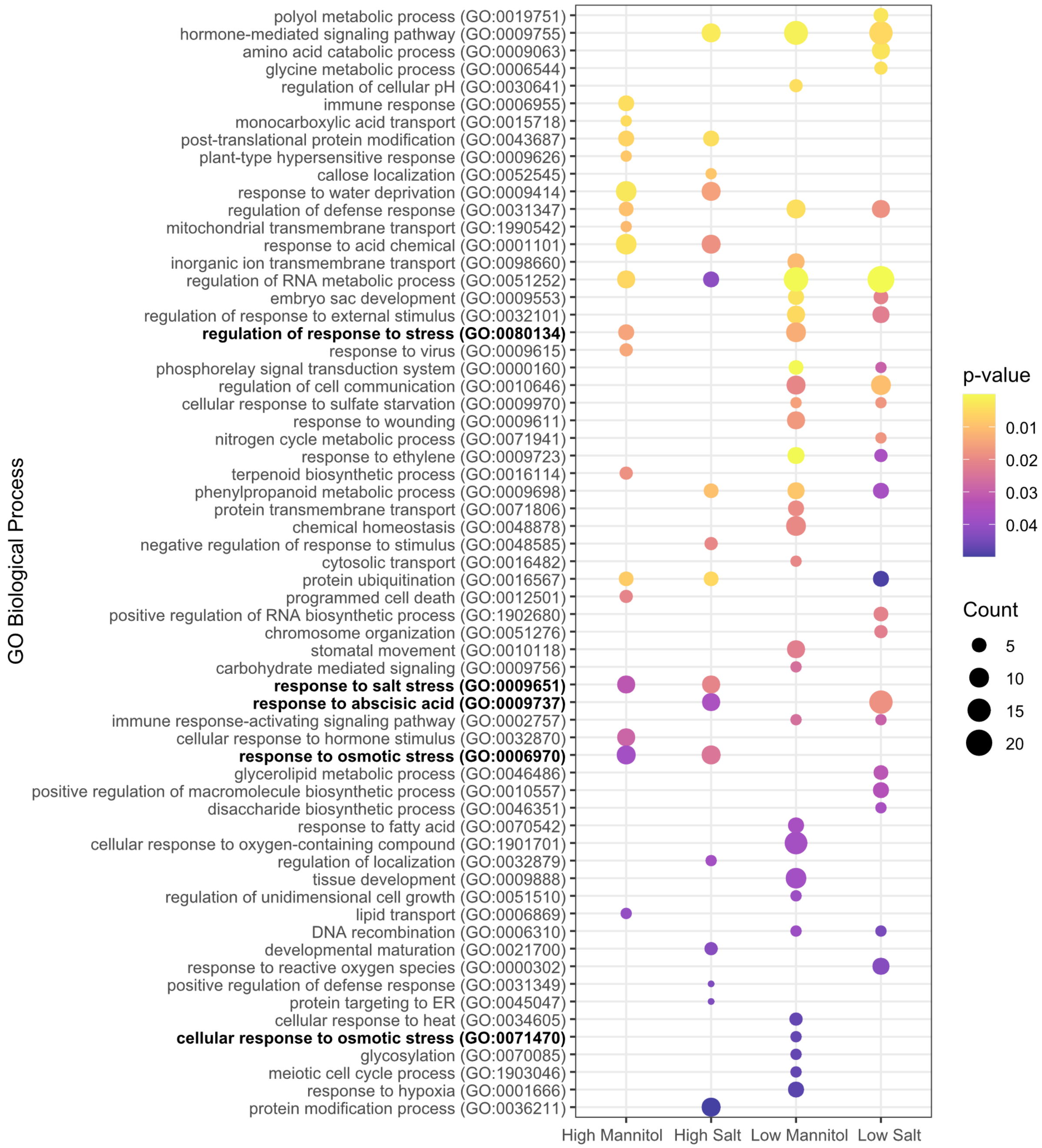


**Supplemental Figure 6. Gene ontology of salt and mannitol stress data specific to AHA-tagged enrichments compared to previous whole proteome analysis.** Gene ontology of biological processes (Benjamini-Hochberg parent-child union, $p < 0.05$) for 'high stress' (300mM mannitol and 150mM salt) and 'low stress' (100mM mannitol and 50mM salt) conditions. All proteins enriched in a stress condition over unlabelled controls in AHA enrichments but not previously detected as significantly changing proteins in the whole proteome analysis of roots or shoots under the same stressor by Rodriguez Gallo et al. (2023) were included as the foreground, with a background of all quantified proteins in the AHA enrichments. Bolded are terms directly related to stress response.
